# Ecophysiological traits of a clonal grass in its climate change response

**DOI:** 10.1101/864827

**Authors:** Veronika Kosová, Tomáš Hájek, Věroslava Hadincová, Zuzana Munzbergova

## Abstract

**Background:** Understanding the ability of species to respond to climate change is essential for prediction of their future distribution. When migration is not adequate, reaction via phenotypic plasticity or genetic adaptation is necessary. While many studies investigated the importance of plasticity and genetic differentiation (plant origin) in growth related traits, we know less about differentiation in ecophysiological traits. In addition, the existing studies looking at plant physiology usually do not estimate the consequences of these physiological changes for species performance.

**Methods:** We used a clonal grass *Festuca rubra* originating from localities representing factorially crossed gradients of temperatures and precipitations. We cultivated the plants in growth chambers set to simulate temperature and moisture regime in the four most extreme localities. We measured net photosynthetic rate, chlorophyll fluorescence, SLA, osmotic potential, stomatal density and stomatal length as range of ecophysiological traits and tested their relationship to plant fitness measured as ramet number and biomass.

**Key results:** We found strong phenotypic plasticity in photosynthetic traits and genetic differentiation in stomatal traits. In most traits, the effects of temperature interacted with the effects of moisture. The relationship between the ecophysiological and fitness-related traits was significant but weak.

**Conclusions:** Ecophysiological response of *Festuca rubra* to climate change is driven by phenotypic plasticity as well as by genetic differentiation indicating potential ability of the populations to adapt to new climatic conditions. The changes in ecophysiological traits translate into plant fitness even though other unmeasured factors also play an important role in fitness determination. Inclusion of species ecophysiology into studies of species adaptation to climate can still increase our ability to understand how species may respond to novel conditions.

## Introduction

Global climate change is underway, and it is predicted to further increase temperature and change precipitation regime (IPCC 2014). Many plant species will thus face novel conditions, which will lead to strong selective pressure on their populations. For understanding species future fitness and distribution, it is necessary to understand the mechanisms which enable the plant species to respond to changing climate (Jump and Penuelas 2005) and identify traits allowing the species to maintain their fitness under novel climate (Gienapp et al. 2008).

Mechanisms, which allow the plants to respond to changing climate, include migration, phenotypic plasticity and genetic adaptation (Anderson et al. 2012a). Migration is well known and documented strategy (Davis and Shaw 2001, Ackerly 2003, Shaw and Etterson 2012). However, for plant species, migration is probably too slow given the contemporary rate of climate change, and therefore the populations will have to face novel conditions *in situ* (Huntley 1991, Davis and Shaw 2001, Gienapp et al. 2008). Phenotypic plasticity, i.e. the ability of a genotype to produce different phenotypes under different environmental conditions, can provide plants better chance to adjust to rapid environmental fluctuations and give them time for further genetic adaptation (Jump and Penuelas 2005, Ghalambor et al. 2007, Nicotra et al. 2010, Valladares et al. 2014). Although phenotypic plasticity could help plants to respond to novel conditions in the long term, it can operate only with the existing genetic variability of the species (Anderson et al. 2012b, Becklin et al. 2016). Genetic adaptations are thus crucial for long-term species response to novel climate.

Genetic adaptation to novel conditions occurs when the species shows sufficient genetic variability which enables selection of genotypes capable to cope with the novel climate (Nicotra et al. 2010, Franks et al. 2014). As a result of such a selection, plants from different populations will become differentiated from each other. The selection may also lead to local adaptation of the plants to their specific environment (Kawecki and Ebert 2004). Existence of genetic differentiation and/or local adaptation provides information about ability of the populations to adapt to their home climate in the past. In combination with sufficient genetic variability and adaptive phenotypic plasticity, it may indicate that the plants will be able be to adapt also to the changed conditions in the future (Gonzalo-Turpin and Hazard 2009).

One of the possibilities to understand species ability to respond to novel climates is to measure key traits, which are under selection under different climatic conditions. Range of previous studies exploring species ability to respond to novel climate focused on differences in plant growth or flowering time, e.g. (Inouye 2008, Gonzalo-Turpin and Hazard 2009, Vitasse et al. 2014, Munzbergova et al. 2017). Nevertheless, climate change causes that plants are increasingly exposed to conditions on their physiological limits (Shaw and Etterson 2012). Proper physiological adaptations are thus crucial for maintaining plant fitness (Becklin et al. 2016) and ecophysiological traits could thus play crucial role in adaptation to changing climate. For example, increased photosynthetic rate or better water extraction in dry environment via osmotic adjustment and changes in stomatal characteristic may increase plant growth and thus improve plant fitness under different climates (Yeh et al. 2012, Benomar et al. 2016). Studies exploring the role of ecophysiological traits for species response to changing climate emerged only in the last decade (8 studies in the review of (Franks et al. 2014), then e.g. (Souther et al. 2012, Zhang et al. 2012, Pratt and Mooney 2013, Sultan et al. 2013, Anderson and Gezon 2015, Knutzen et al. 2015, Benomar et al. 2016, Acuna-Rodriguez et al. 2017, Hamann et al. 2018, Prud’homme et al. 2018).

The studies exploring ecophysiological responses of plants to climate showed that ecophysiological traits are often highly plastic. Approximately in half of the cases, also genetic differentiation between populations has been detected. Some traits (e.g. stomatal traits) tend to show significant genetic differentiation, but no plasticity. Despite the accumulated knowledge on plasticity and genetic differentiation of the ecophysiological traits, it is not clear how the changes in these traits affect the ability of plants to maintain their fitness under novel conditions (Becklin et al. 2016). Studies measuring both ecophysiological and fitness related traits in the same system and explicitly looking for their relationships, are, however, still sparse (e.g. (Pavlikova et al. 2017, Munzbergova and Haisel 2019)). Moreover, very few of them looked at such a relationship while exploring species response to novel climates e.g. (Vogan and Maherali 2014, Carlson et al. 2016, Hamann et al. 2016, Ramirez-Valiente et al. 2017).

All studies focusing on phenotypic plasticity vs. genetic differentiation in ecophysiological traits described above used plant populations growing either along temperature or precipitation gradients, and nothing is known on their interactions. Nevertheless, the interaction between these factors is also very important as shown in a range of studies exploring patterns of plant growth (Meineri et al. 2014, Moran et al. 2016, Munzbergova et al. 2017).

Here, we used unique system of localities with factorially crossed gradients of temperature and precipitation located in western Norway, previously used in several other studies, e.g. (Meineri et al. 2013, Meineri et al. 2014, Klanderud et al. 2015, Munzbergova et al. 2017). The system provides a unique possibility to explore the importance of independent as well as combined effects of temperature and precipitation on different ecophysiological traits and thus to evaluate the importance of including interactions between different climatic factors into models attempting to predict future species response to changing climate (Butof et al. 2012, Lu et al. 2016, Wu et al. 2016).

To study species ecophysiological responses to climate, we measured 6 different ecophysiological traits expected to be important for maintaining species fitness (Benomar et al. 2016). (i) Net photosynthetic rate (P_N_) is usually positively correlated with plant production (Peng et al. 1991). It could change with temperature (Yamori et al. 2014) and also could be affected by soil moisture via stomatal regulation (Azhar et al. 2014). (ii) Maximum photosystem II efficiency (Fv/Fm) reflecting species response to various kinds of stresses in reaction to different environmental conditions such as drought or temperature stress (Maricle and Adler 2011). (iii) Osmotic potential is key drought resistance trait with high level of plasticity (Bartlett et al. 2014). (iv) Specific leaf area (SLA) is often positively correlated with plant production (Garnier et al. 2001) and nitrogen concentration (Wright et al. 2004) and may be indicative of trade-off between rapid biomass production (high SLA) and efficient nutrient conservation (low SLA) (Giuliani et al. 2014). (v) Stomatal density and (vi) stomatal length regulate gaseous exchange (Bussotti et al. 2005) and thus are crucial for photosynthesis (Beerling and Chaloner 1993). All these traits already showed high level of phenotypic plasticity as well as genetic differentiation between populations in other systems (Bussotti et al. 2005, Agrawal et al. 2008, De Frenne et al. 2011, Reinhardt et al. 2011, Robakowski et al. 2012, Taylor et al. 2012, Azhar et al. 2014, Montoro and Sadras 2014, Xu et al. 2014, Yamori et al. 2014, Psidova et al. 2015, Carlson et al. 2016, Song et al. 2016).

By linking our ecophysiological measurements with previously published data on plant fitness (Munzbergova et al. 2017), we also explored whether and how the changes in plant ecophysiology are translated into plant fitness. We used wide-spread clonal grass, *Festuca rubra*, as a model for the study. *F. rubra* is important dominant species of many grassland systems and clonal reproduction represents very important strategy for many grassland species. By exploring this species, we are likely to understand responses of the major components of many ecosystems especially in higher elevations. Specifically, we aim to answer the following questions: (1) What is the relative importance of phenotypic plasticity, genetic differentiation and their interactions for the ecophysiological traits? (2) Can we identify signs of local adaptation, i.e. do plants perform better in their home conditions? (3) How do the plants respond to specific changes in climatic conditions and what is the relative importance of change in temperature and precipitation and their interaction? (4) Is there any relationship between ecophysiological traits and fitness, i.e. does selection play a role in the ecophysiological traits, and does this relationship depend on climate?

We hypothesized that: (1) both genetic differences and plasticity will contribute to variation in the ecophysiological trait response to climatic gradients and their change, with interactions between conditions of plant origin and of its actual growth indicating that the ability to respond to climate change varies across the species’ climatic niche as previously shown for growth and fitness related traits in the same system (Munzbergova et al. 2017). (2) Climate of origin exerts strong selection pressure on the local populations. Therefore, we expect that mainly in the ecophysiological traits showing strong genetic differentiation between populations, local adaptation will be detected. (3) All plants will profit from transplantation to warmer and wetter conditions, i.e. from the projected climate change in the region, as such conditions are likely more favourable for the species (Munzbergova et al. 2017). (4) The physiological traits will be related to plant fitness and the association will be the strongest in most favourable conditions, likely represented by the warm-wet conditions.

## Methods

### Plant material

*Festuca rubra* L. is a common perennial grass species of temperate grasslands in Europe. *F. rubra* used in the experiment is a widespread hexaploid type from the *F. rubra* complex (Šurinová et al., in press.). The species grows across a wide climatic gradient in grasslands, both as a dominant with only few other species and also as a minor component of species rich stands. It reproduces by seeds as well as vegetatively producing both intravaginal and extravaginal tillers on rhizomes (Münzbergová et al. 2017). Clones of *F. rubra* are characterized by considerable genetic variability and plasticity in response to environmental factors (Skalova et al. 1997, Herben et al. 2001, Münzbergová et al. 2017).

### Study sites

The plant material was collected in western Norway at localities distributed along a natural climate grid described in detail by ((Meineri et al. 2012, Meineri et al. 2013, Meineri et al. 2014, Klanderud et al. 2015, Vandvik et al. 2016)). The grid consists of 12 sites, arranged along four levels of mean annual precipitation (ca. 600, 1300, 2000, and 2700 mm) and three levels of summer temperature (means of the four warmest months (6.5 °C – alpine, 8.5 °C – subalpine and 10.5 °C – boreal) (Fig. 1). The communities are grazed intermediate-rich meadows with comparable slopes, grazing regime and bedrock (Meineri et al. 2014, Münzbergová et al. 2017).

**Fig. 1.**
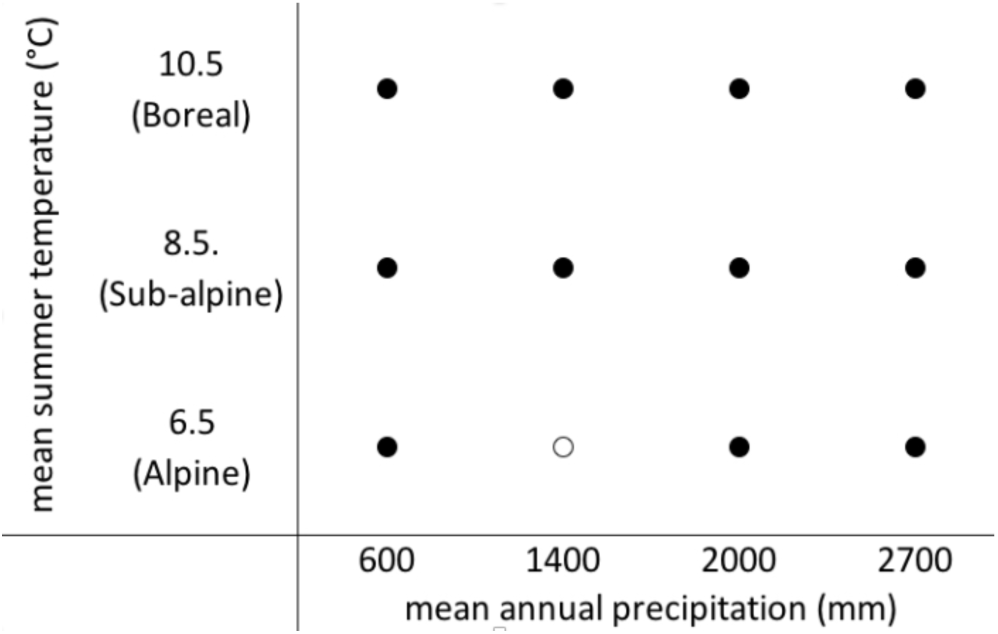
Position of original localities within SeedClim climate grid along the temperature and precipitation gradient. The black dots represent all the localities at which *Festuca rubra* was collected, the white dot shows the locality, from which *Festuca rubra* was not available.

### Experimental design

We used the same plant material as in a previous experiment (Münzbergová et al. 2017), which contained 25 genotypes of *F. rubra* hexaploids originating from each locality except alpine locality with 1400 mm of precipitation (Fig. 1), where no *F. rubra* hexaploids were available. Each genotype was vegetatively reproduced to gain four ramets from each genotype (to have 4 × 25 ramets from each population). The ramets were planted into pots 5 × 5 × 8.5 cm into a mixture of common garden soil and sand in 1:2 ratio. The pots were divided into 4 groups so that one ramet from each genotype was represented in each group. Thus, in each group there were 11 populations × 25 genotypes, i.e. 275 ramets resulting in 1100 ramets in total. Pots within each group were fully randomized and moved to a growth chamber. We randomized the position of the pots in the growth chambers monthly. The plants grew in the growth chambers for 176 days.

The plants were cultivated in four growth chambers (Vötch 1014) under conditions simulating the field situation in the spring (second half of April - second half of June). The regimes were derived from temperature course in the four extreme localities (wettest and driest combined with warmest and coldest) but respecting technical possibilities of the growth chambers and thus avoiding night frosts. The temperatures in the growth chambers differed between the cold and warm treatments and changed over the growing season following the course of temperature at the natural localities. The course of temperatures during the growth chamber experiment is shown in supplementary material (Table S2). For more details about collecting, identifying, planting the plants and exact environmental parameters of the growth chambers see (Münzbergová et al. 2017), where the same plant material was used.

### Plant traits

The plants grew in the growth chambers for 176 days. After this time, we measured the ecophysiological photosynthesis-related traits: net photosynthetic rate (P_N_), maximum efficiency of photosystem II photochemistry (parameter Fv/Fm), specific leaf area (SLA), osmotic potential, stomatal density and stomatal length. At the same time, growth-related traits serving as proxies of fitness (number of ramets, belowground biomass and aboveground biomass) were measured for a previous study (Münzbergová et al. 2017). Here, we used these growth-related traits to assess phenotypic selection of the ecophysiological traits (see below).

Net photosynthetic rate (P_N_), defined as the total rate of photosynthetic CO_2_ fixation, was measured using an infra-red gas analyzer LI-6400 (LI-COR, Lincoln, Nebraska, USA). The measurements were done at 25° C ±2° C and minimum 55% of relative humidity. LI-COR was set at light intensity of 1600 μmol m^−2^ s^−1^, air flow rate 300 µmol s^−1^ and CO_2_ concentration of 400 ppm. Each plant was measured for at least 25 minutes to ensure that the stomatal conductance is stabilized. Because these measurements are very time consuming, they were done using randomly chosen 10 plants originating from each of the most extreme localities: wettest and driest combined with warmest and coldest, i.e. 40 plants in each growth chamber, i.e. 160 plants in total. Final P_N_ is expressed per unit area (mm^2^). To verify that time of day had no effect on measurement, several plants were pre-experimentally measured three times during a day - in the morning, at midday and in the afternoon.

SLA was calculated as a ratio of leaf area (mm^2^) to dry mass (mg). Leaf area was calculated on scanned folded leaves (as *Festuca* leaves are naturally folded and are often hard to unfold) multiplied by two to acquire real area of the leaves. We used the same reduced sample as for P_N_ measurements. This is because the leaf area data primarily served for calculating the P_N_, as described above.

Maximum photosystem II (PSII) efficiency - Fv/Fm was measured using FluorPen FP-100 MAX/USB (Photon System Instruments, Czechia) in dark-acclimated (1 h) plants (Maxwell and Johnson 2000).

Osmotic potential was determined psychrometrically from leaf sap using dew point microvoltmeter Wescor HR-33T (Wescor, Inc., USA) with C-52 sample chambers.

Stomatal density and stomatal length were measured on epidermal imprints made by applying nail polish on a fresh leaf surface and gently peeled from the surface using transparent adhesive tape after polish hardening (Gitz et Baker 2009). The impressions were mounted on microscope slides with the transparent adhesive tape. Stomatal density was counted as an average of three non-overlapping areas (each 500 × 500 µm). Each counting area was in contact with margin of the leaf. From each of these three areas, three stomata were randomly chosen, and their length was measured, resulting in 9 measured stomata per sample. Stomatal density and stomatal length were measured on samples of all localities, but we randomly selected only 10 plants from each locality. The selection of plants originating from the most extreme localities was kept according to selection for measurement of net photosynthetic rate and SLA.

## Data analysis

All tests were done using mixed effect models with clone identity as a random factor. If it was necessary, data were transformed to fit the assumptions of normality and homogeneity of variance. For all the dependent variables, the tests assume Gaussian distribution of the dependent variable, except for Fv/Fm assuming Gamma distribution. Significance was derived from LR-tests (likelihood ratio tests of goodness of fit (Bolker 2009)). Mixed effect models were processed with R (version 3.3.2) using lme4 package (Bates et al. 2015).

For all tests, we used sequential Bonferroni correction (Rice 1989) as we used 6 different traits measured on the same experimental plants. Because the necessity to use the correction is ambiguous (as described in (Munzbergova et al. 2017)), we decided to illustrate results both with and without this correction.

### Effects of target and original conditions

Data analyses used the same strategy as a previous study on growth and foraging related traits of the same plants in the same system (Munzbergova et al. 2017). We tested the effect of original conditions, target conditions (i.e. conditions of cultivation in the growth chambers) and their interactions on all the ecophysiological traits. For this purpose, we coded the temperature of origin according to the mean temperature of the four warmest months at the original localities as 6.5, 8.5 and 10.5 °C and the precipitations of origin with respect to the mean annual precipitation at the original localities as 600, 1300, 2000 and 2700 mm. The target conditions in the growth chambers were coded in the same way as 6.5 and 10.5 °C for temperature and 600 and 2700 mm for precipitation regime. The significant effect of target conditions indicates plasticity in the traits of the plants, the effect of origin indicates genetic differences between plant populations originating from different climatic conditions and the interaction between target and origin indicates genetic differentiation in plasticity. For net photosynthetic rate, additional analysis with time of measurement as covariate was done to check the influence of measuring time on the results. The time had no significant effect on the results, thus it is not used in further analyses.

The analyses described above were done on different numbers of observations, which depended on number of measured samples for each trait. To assess the effect of sample size on the results, we made additional analyses with reduced numbers of samples (160) according to the traits with the lowest number of samples (net photosynthetic rate and SLA). These analyses are shown in supplementary materials (Table S1). Relative importance of local and target conditions was determined as the proportion of total deviance explained in the data.

### Local adaptation

To study signatures of local adaptation, local vs. foreign criterion was used (Kawecki and Ebert 2004). This criterion presume that the plants prosper better in their home conditions compared to foreign conditions. For this test, the plants grown in the same temperature or moisture as they had in original localities were marked by 1. Plants that grew in foreign conditions were marked by 0. Here, we tested the effect of local vs. foreign temperature, moisture and their interaction on all the ecophysiological traits.

### Directionality in environmental change responses

Then, we explored how the plants respond to the directionality of climate change between the original localities and the growth chambers. To assess the directionality of climate change, we subtracted the codes for original localities from codes for target conditions. It resulted into new codes (−2100 mm to 2100 mm for moisture and −4 °C to 4°C for temperature). The negative values indicate transplantation to colder or drier conditions in the growth chamber, while the positive values indicate transplantation to warmer or wetter conditions. Zero indicates plants grown in their home conditions. We tested the effect of differences in temperature, moisture and their interactions.

### Relationship between ecophysiological traits and relative fitness

We tested phenotypic selection of ecophysiological traits by regressing them against traits representing plant fitness. Specifically, we used three growth-related traits (number of ramets and aboveground and belowground biomass) instead of classically used fitness traits as number of seeds or number of flower-buds as fitness measure. This is because the growth-related traits are probably more suitable proxies of fitness for long-lived plant species with predominant clonal reproduction as explained in (Munzbergova et al. 2017). Specifically, we used standardized linear (directional) selection gradients (β), calculated by means of multiple linear regressions (i.e. the partial regression coefficient from the multiple regression of relative fitness on all standardized ecophysiological traits) (Lande and Arnold 1983). Selection gradient was calculated on reduced sample size (n=160) according to the number of observations for net photosynthetic rate and SLA. We also estimated univariate selection differential (*S*), calculated as linear regression between each ecophysiological and fitness traits, which include both direct selection and indirect selection caused by trait correlations (Lande and Arnold 1983). Selection differentials were based on full sample (i.e. n=160 for net photosynthetic rate and SLA, n=440 for stomatal density and length, n=1100 for osmotic potential and Fv/Fm). For both type of tests, we used number of ramets, aboveground biomass and belowground biomass as proxies of fitness (Munzbergova et al. 2017, Younginger et al. 2017) and we calculated relative fitness by dividing each value by the mean of the values of the corresponding growth chamber. Ecophysiological traits were standardized to a mean of zero and a variance of one. Population of origin was used as a covariate in these tests.

## Results

### Effect of target conditions

Target conditions (conditions in the growth chambers) significantly affected all the measured traits (Fig. 2, Table 1). The strongest effect of target conditions was found in net photosynthetic rate (P_N_), specific leaf area (SLA), maximum photosystem II efficiency (Fv/Fm) and osmotic potential explaining more than 70% of explained deviance in these traits (Fig. 2). Specifically, target temperature and moisture affected P_N_, Fv/Fm, SLA and stomatal length. Stomatal density did not respond to their separate effects, but only to their interaction. Osmotic potential was not affected by target moisture, but it was affected by the target temperature (Table 1). In general, the plants in dry growth chambers had higher values of P_N_, Fv/Fm, SLA, and stomatal length (Fig. S1A, B, C, F). The plants in warm growth chambers had higher Fv/Fm, SLA, osmotic potential, and stomatal length. Only P_N_ was higher in cold growth chambers.

**Fig. 2.**
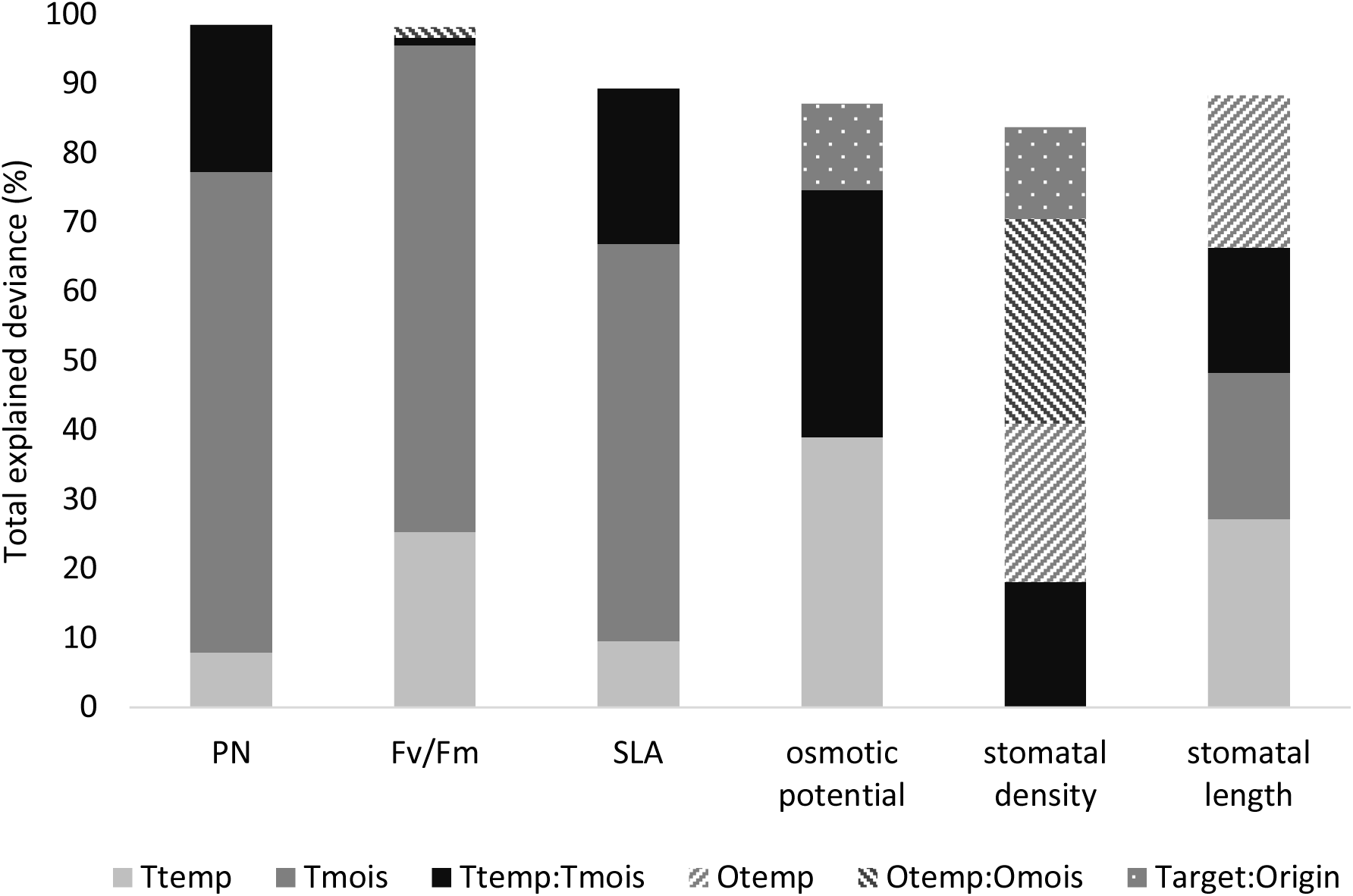
Relative importance of original and target conditions and their interaction in all measured traits. Percentage shows total explained deviance by the model for significant variables. Original moisture is not shown as it had no significant effect on any of the traits.

We found significant interactions between target temperature and moisture in all traits (Table 1). Specifically, P_N_ and SLA increased with increasing temperature in wet growth chambers, while decreased with temperature in dry growth chambers. Fv/Fm also increased with increasing temperature and the increase was stronger in wet growth chambers (Fig. S1A, C). Osmotic potential increased with increasing temperature in wet growth chambers and did not change with temperature in dry growth chambers (Fig. S1D). Stomata density and length were constant in wet growth chambers, while in dry growth chambers stomatal density decreased and stomatal size increased with temperature (Fig. S1E, F). All target conditions effects were significant after applying the Bonferroni correction, except the effect of target moisture on stomatal length (Table 1). In the analyses with reduced sample size (Table S1), there was additional significant effect of moisture on osmotic potential. On the other hand, previously significant effect of interaction between target temperature and moisture on osmotic potential, stomatal density and stomatal length became non-significant (Table 1, Table S1).

### Effect of original conditions

Original climate significantly affected Fv/Fm, stomatal density, and stomatal length (Table 1). The strongest effect of original conditions was found in stomatal density, stomatal length and osmotic potential (Fig. 2). Specifically, increasing original temperature negatively affected stomatal density (Fig. S1E) and positively stomata length (Fig. S1F). Original moisture did not have any significant effect (Table 1). Interaction between original temperature and moisture affected Fv/Fm and stomatal density (Fig. 2, Table 1). Stomatal density as well as Fv/Fm increased with increasing moisture in cold conditions, while it decreased in warm conditions (Fig. S1B, E). After applying the Bonferroni correction, the effect of original conditions remained significant for stomatal density only. The significant effects of original conditions remained the same in analyses on the reduced number of samples (Table S1).

### Interaction between target and original conditions

Two significant interactions between original and target conditions were found (Table 1), but the strength of these interactions was lower than the strength of the main effects (Fig. 2). Specifically, osmotic potential was affected by interaction between original and target moisture, and stomatal length by interaction between original and target temperature (Fig. 2, Table 1). Osmotic potential of plants originating from drier localities decreased when shifted to wet conditions. In contrast, it decreased in plants coming from wetter localities with shifting to drier conditions (Fig. S1D). Stomatal density of plants coming from cold climate differed more from other plants in warm growth chambers than in cold growth chambers (Fig. S1E). Nevertheless, none of these effects remained significant after applying Bonferroni correction (Table 1). Results of analyses with reduced sample size were different from full-size analyses (Table 1, Table S1). In the reduced analyses, interaction between original and target temperature had no effect on stomatal length but on stomatal density. Osmotic potential was not affected by interaction between original and target moisture. In addition, there was significant interaction between original temperature and target moisture as well as between original moisture and target temperature on Fv/Fm (Table S1).

### Signs of local adaptation

Difference between local and foreign conditions was significant for osmotic potential and for stomatal length. Osmotic potential depended on local/ foreign moisture, and stomatal length was affected by interaction between local/ foreign temperature and moisture (Table S3). Specifically, plants growing in foreign moisture had lower osmotic potential than plants growing in local moisture (Fig. S3A). Plants growing in local temperature and moisture had shorter stomata than in other combinations (Fig. S3B). Difference between local and foreign conditions was not significant for any trait after applying Bonferroni correction (Table S3).

### Directionality in environmental change responses

In all traits except stomatal length, significant effect of changing conditions was detected. Specifically, raising temperature had significant positive effect on SLA, Fv/Fm, and osmotic potential, but negative effect on P_N_ (Fig. S2A, B, C). Rising moisture had significant negative effect on P_N_, Fv/Fm, SLA and stomatal density (Table 2, Fig. S2A, B, C, D).

Interaction between change in temperature and moisture was significant for P_N_, SLA and osmotic potential (Table 2). In case of P_N_, with shifting to drier conditions the values were generally higher and decreased with increasing temperature, while with shifting to wetter conditions the values were generally lower and independent of temperature (Fig. S2A). SLA decreased with the temperature when moisture was reduced, while SLA increased with increased moisture in increased temperature (Fig. S1C). Osmotic potential increased with temperature in wetter climate, while temperature did not play any role in drier climate (Fig. S1D). All effects were still significant after applying Bonferroni correction, except effect of difference in moisture on stomatal density (Table 2).

### Relationship between ecophysiological traits and relative fitness

The phenotypic selection analysis showed several significant selection gradients, suggesting relationship between ecophysiological traits and fitness-related traits. The results strongly differ between traits as well as between specific target conditions (Table 3). Overall, the directional selection expressed using selection gradients was very weak in wet target conditions, where we only found a significant negative relationship between P_N_ and belowground biomass, and positive relationship between stomatal length and belowground biomass. In dry-cold conditions, we found significant negative relationship between P_N_ and aboveground biomass. Warm-dry conditions were found to be the most selective. Here we detected significant positive relationship of SLA, osmotic potential and stomatal density with ramets number, and significant negative relationship between osmotic potential and aboveground biomass. Only 3 out of 9 results were still significant after Bonferroni correction (relationship between SLA and osmotic potential and ramet number in warm and dry conditions, and relationship between stomatal length and belowground biomass in cold and dry environment, Table 3).

In contrast to directional selection gradients, 10 additional relationships became significant when calculating selection differentials (Table 3). Mainly, these are the relationships between fitness proxies and osmotic potential or Fv/Fm (8 out of 10). In contrast to directional selection gradients, osmotic potential was positively selected in all growth conditions when estimating selection as selection differentials. Fv/Fm was positively related to belowground biomass in cold-wet conditions but negatively related to number of ramets and aboveground biomass in warm conditions. Relationship between stomatal length and belowground biomass in cold-dry conditions became non-significant as well as the relationship between number of ramets and stomatal density in warm-dry conditions. In contrast, stomatal density relationship with number of ramets and aboveground biomass became significant in dry environments. Selection differentials of net photosynthetic rate as well as specific leaf area remained consistent with directional selection gradients. All these 15 significant results except two (number of ramets with osmotic potential and SLA) became non-significant after Bonferroni correction (Table 3).

## Discussion

The ecophysiological traits of our studied species, *F. rubra*, showed high degree of phenotypic plasticity as well as genetic differentiation in response to temperature and moisture. It indicates that both mechanisms – genetic differentiation and particularly phenotypic plasticity - are important for the ability of this species to cope with changing climate. This is in line with other studies exploring genetic and plastic responses using ecophysiological traits (e.g. (Souther et al. 2012, Zhang et al. 2012, Pratt and Mooney 2013, Sultan et al. 2013, Anderson and Gezon 2015, Knutzen et al. 2015, Benomar et al. 2016). Nevertheless, the relative importance of phenotypic plasticity and genetic differentiation in given traits strongly differs among species as well as among studied climatic factors. All the previous studies used populations across either temperature or precipitation gradient for simulation of climate change. Our study, however, demonstrated that also the interaction between temperature and moisture is very important.

We also tested the importance of ecophysiological traits in our study by relating them to plant fitness. We found many significant relationships indicating that changes in the ecophysiological traits may affect plant fitness and thus species ability to respond to climate change. However, it is important to keep in mind that most of the significant relationship were rather weak, suggesting that the strength of the relationships needs to be carefully considered when interpreting the results. Nevertheless, the strengths of the relationships correspond to the strengths detected previously (e.g. (Sherrard and Maherali 2006, Agrawal et al. 2008, Lopez-Gallego and O’Neil 2014, Hamann et al. 2016, Ramirez-Valiente et al. 2017), but see e.g. (Cano et al. 2008, McIntyre and Strauss 2014, Sherrard et al. 2015, Carlson et al. 2016)). This suggest that other processes also play an important role in fitness determination.

### Phenotypic plasticity

Phenotypic plasticity, determined by significant effect of target conditions, was shown in all traits. However, net photosynthetic rate, SLA, Fv/Fm and osmotic potential were more plastic than stomatal density and stomatal length. High plasticity of photosynthetic parameters (net photosynthetic rate, Fv/Fm) was consistently demonstrated also in previous studies (Souther et al. 2012, Yamori et al. 2014, Benomar et al. 2016). In contrast, plasticity of SLA, osmotic potential and stomatal traits differs between study systems and species ((Ramirez-Valiente et al. 2010, Avolio and Smith 2013, Pratt and Mooney 2013, Knutzen et al. 2015).

Low net photosynthetic rate, SLA and Fv/Fm reflect plant stress (Shi et al. 2010, Ashraf and Harris 2013). According to these traits, the plants were the most stressed in cold-wet target conditions, and the least stressed in dry and particularly in the cold-dry conditions. All these traits were affected by temperature, moisture and their interaction, but the main driver of the differences between plants growing in the different growth chambers was moisture. Lower level of stress in dry chambers contrasts to other studies (e.g. (Ramirez-Valiente et al. 2010, Pratt and Mooney 2013, Sultan et al. 2013) indicating that plants are more stressed under dry conditions as judged based on P_N_, SLA and Fv/Fm. However, wet conditions in our experiment represent high annual precipitations of 2700 mm and our dry conditions represent annual precipitations of 600 mm. Our dry conditions thus correspond to what would be considered wet in many regions. It is possible that, especially in combination with low temperature, plants under high and permanent soil moisture could face hypoxia, suggested by values of net photosynthetic rate, SLA and Fv/Fm, and accumulation of protective carotenoids (detected as visible red pigmentation of leaves in our experiment), which could help in protecting the plants against hypoxia (Ougham et al. 2005, Kawasaki et al. 2015). Thus, the plant reaction seems to be in line with ecology of the species.

The effect of temperature on Fv/Fm, SLA and net photosynthetic rate slightly differed between these traits. Fv/Fm increased with rising temperature indicating lower light-related stress in warmer conditions (Maricle and Adler 2011). SLA was higher in warmer conditions, which corresponds to other studies transplanting plants along latitudinal gradients (Gonzalo-Turpin and Hazard 2009, Benomar et al. 2016). But in case of net photosynthetic rate, the reaction to warm conditions was negative, which was primarily driven by the highest values in cold-dry growth chamber. The difference between net photosynthetic rates in warm and cold growth chambers was, however, negligible in contrast to difference between wet and dry growth chambers. It indicates that, despite the significant effect of temperature, the driver of this difference is mainly moisture.

Low values of osmotic potential indicate plant acclimatization to dry conditions (Bartlett et al. 2014). Here, osmotic potential was not affected by target moisture (as the main effect). The absence of effect of moisture on osmotic potential corresponds with the study of (Knutzen et al. 2015), but they used relatively short moisture gradient. We suggest that target moisture in our study did not affect osmotic potential because the plants never suffered from severe drought in the experiment. Surprisingly, osmotic potential was significantly affected by temperature and by interaction between temperature and moisture. The lowest values of osmotic potential were detected in the cold growth chambers. It could mean that the plants tried to prevent cold stress through accumulating of osmotic solutes in their leaf cells. Nevertheless, maximum difference between measured values was 0.1 MPa, which probably could have very small effect on plant physiology.

Stomatal density and stomatal length showed significant phenotypic plasticity, but especially in case of stomatal density the plasticity was very low. These results differ from previous studies, which used temperature gradient and demonstrated plasticity in stomatal density, but not in stomatal length (Zhang et al. 2012, Anderson and Gezon 2015). Thus, it seems that degree of plasticity of stomatal traits is species-specific.

Altogether, the data on phenotypic plasticity suggest that the plants are able to react to different climatic conditions plastically without significant signs of their damage. Overall, the plants performed the best in dry and warm conditions, but the main driver of plant performance was moisture.

### Genetic differentiation

Only three out of the six ecophysiological traits were affected by climate of origin, namely Fv/Fm, stomatal density and stomatal length, indicating genetic differentiation between populations. In case of Fv/Fm, the effect of original conditions (interaction between original temperature and moisture) was negligible as compared to the effect of target conditions. Thus, only stomatal traits showed substantial genetic differentiation between populations.

Stomata traits were influenced by original temperature and by interaction between original temperature and moisture. We could expect that plants originating from warmer and/or drier conditions have more abundant but smaller stomata (Xu and Zhou 2008, Yan et al. 2017). This prediction was confirmed only partly in the warm localities, where stomatal density slightly increased with decreasing precipitations. However, in case of this study, it is necessary to consider specificity of our original localities. Our “dry” localities have, in fact, relatively high annual precipitation and our “warm” localities with mean summer temperature of 10.5°C are less stressful for our model plants than the “cold” localities. Thus, our finding of less numerous and bigger stomata in warm localities could be caused simply because plants in these localities have higher photosynthetic rate than plants originating from colder localities and thus they do not need to invest to more numerous stomata to try to compensate photosynthetic rate by higher intake of CO_2_. Similarly, plants could produce the highest stomata density at coldest-wettest locality as they have enough water there and they need to maximize effectiveness of photosynthesis. In case of temperature, the effect could be also enhanced by different altitudes of origin.

Stomatal density can be higher in colder conditions as it increases with decreasing CO_2_ concentration (Woodward et al. 2002), i.e. with increasing altitude and thus decreasing temperature. Stomatal density can also increase because of increased solar radiation intensity in high altitudes (Yan et al. 2017). Possible importance of other environmental factors for trait differentiation was also shown in a study exploring adaptive differentiation using molecular markers and quantitative traits in the same study system ((Stojanova et al. 2018). Nevertheless, Stojanova et al. (2018) also demonstrated that stomatal density showed (as the only one of many traits) patterns of adaptive differentiation and moreover, the trait also showed strong divergent selection in plasticity. It indicates that the populations originating from different environment have different plasticity (the most plastic populations were those originating from the coldest localities). It could reflect different spatial or temporal heterogeneity of the different environments with the extremely cold conditions being the most heterogenous. Altogether, it indicates that the possibility of adaptation through this trait exists. Nevertheless, specific expression of stomatal traits are probably species and environment-specific because in other studies the effects of genetic differentiation strongly differed between species and study systems (Zhang et al. 2012, Anderson and Gezon 2015, Yan et al. 2017).

### Interaction between target and original conditions

The interaction between target and original conditions represents genetic differences in plasticity between populations (e.g. (Hamann et al. 2016)). These differences were significant for osmotic potential and stomatal density. This interaction indicates that phenotypic plasticity in these traits could be under selection (Thompson 1991) and populations with higher adaptive plasticity (i.e. positive reaction to changing climate (Ghalambor et al. 2007)) could have better chance to adjust to novel conditions. However, the range of the actual values of both osmotic potential and stomatal density was very low and therefore a biologically relevant effect cannot be assumed. Weak interaction between target and original conditions was also found by (Gugger et al. 2015) and (Munzbergova et al. 2017).

### Local adaptation

Despite significant effect of home vs. foreign criterion on osmotic potential and stomatal density, local adaptation was not unambiguously proved. Osmotic potential of plants growing in their home conditions was higher than those growing in foreign condition. It could probably mean that the plants tried to adjust to novel (stressful) condition. Nevertheless, it is important to take in consideration, that differences in the values between home and foreign conditions in the osmotic potential were only approximately 0.03 MPa, which may be too low to have any real effect. We are, however, not aware of any study, which would provide numerical information on the relevance of such a change in the osmotic potential for plant fitness. In case of stomatal density, plants in their home temperature and moisture had smaller stomata than in other combinations, but the mechanism of this effect is unclear. Nevertheless, stomatal density and length showed high level of genetic differentiation and evident differences between populations. It could mean that despite unclear home vs. foreign analysis, the stomatal traits showed dependence on plant origin and thus the potential for local adaptation through these traits exists (Kawecki and Ebert 2004). Our result is in line with (Munzbergova et al. 2017), and the result indicate that the plants are not be constrained by local adaptation as it could be limiting species ability to adapt to novel climate (Aitken and Whitlock 2013).

### Effect of changing climate

The general climatic prediction for Norway suggests increases in both precipitation and temperature over the next century, but high temperature fluctuations with longer drought periods alternated by intense rainfalls are also expected (IPCC 2014). Out of all the measured traits, only net photosynthetic rate, SLA and Fv/Fm have a biologically relevant effect on the response to directionality of climate change, despite the direction of the change had a significant effect on all traits except stomatal length. The increased temperature had overall positive effect on the plants, the effect of increased precipitation was negative. Thus, within these photosynthesis-related traits, it seems that the plants will not benefit from general predicted climate change. However, when the effect of changing climate was studied on growth-related traits (Münzbergová et al. 2017), shift to warmer and wetter conditions was beneficial for the model plants. This inconsistency is probably caused by the fact that the most plastic traits presented here were negatively influenced mainly by increasing precipitation. In Munzbergova et al. (2017), the transplantation to wetter conditions decreased plant height and number of ramets, but this effect was mitigated by rising temperature. Thus, plants are likely to profit from rising temperature, but rising precipitation will more or less inhibit this positive effect.

### Relationship of ecophysiological and fitness traits

Significant relationship between fitness and ecophysiological traits was found for all traits, which indicates that all of them should be under selection. Nevertheless, the selection seems to be very weak. According to (Becklin et al. 2016), ecophysiological traits could be under strong selection caused by climate change and therefore, we expected that plant fitness is strongly dependent on ecophysiological traits and thus the ecophysiological traits are under strong selection. We were unable to clearly confirm this prediction. One of the possible reasons is that despite long climatic gradients in our study system, we lack variance around mean values of temperature and precipitations, which could represent extreme conditions (e.g. extremely high summer temperature in combination with drought). Most studies which demonstrated selection on ecophysiological traits used systems with natural or simulated precipitation gradients where the plants were exposed to extreme drought (e.g. (Vogan and Maherali 2014, Carlson et al. 2016, Quezada et al. 2017)). Nevertheless, their results on strength of selection strongly differ between traits and species. Another possible reason why we found weak relationship could be incorrect selection of fitness proxies. But for the reasons described in methods, we assume that number of ramets and biomass are good fitness proxies in our system.

Despite weak significant results, we found negative relationships of net photosynthetic rate with fitness in two contrasting environments – in warm-wet and cold-dry conditions. Usually, net photosynthetic rate is positively correlated with biomass production (Peng et al. 1991), but they can become decoupled under additional stress that affects assimilate allocation strategies, such as nutrient or water limitation and in our case also likely water access. Fv/Fm was positively selected in cold-wet conditions and negatively selected in warm environments denoting light-induced stress under potentially hypoxic soil conditions. Stomatal density had no effect on fitness in wet conditions where the plants could maintain maximum stomatal conductance. But in dry environments, stomatal density was positively selected in cold conditions and negatively in warm conditions, perhaps reflecting the demand for CO_2_ supply under contrasting photosynthetic rates affected by hypoxia mentioned above, rather than controlling the CO_2_ supply. The selection for longer stomata in cold environment independent of moisture may indicate optimum water supply as fewer bigger stomata are expected to be less costly than numerous small stomata (Franks et al. 2009).

For osmotic potential, we found negative selection in warm-dry conditions and positive selection in the remaining environments. The expected, but only slightly negative, selection in warm-dry conditions represents a typical response to drier environment (Bartlett et al. 2014). In the other climatic conditions, the lack of any water deficit suppressed any differentiation in osmotic potential.

We found positive selection for SLA in warm-dry environment. The selection for higher SLA suggests optimum growth conditions and indicates that the higher SLA promoted resource allocation to vegetative spread.

## Conclusions

Our main aim was to understand the importance of ecophysiological traits in response of *F. rubra* to climate change. This study showed high level of phenotypic plasticity in all the traits with the highest levels in photosynthesis-related traits. Populations originating from different climatic conditions were also genetically differentiated in stomatal traits. Thus, the importance of plasticity and genetic differentiation strongly differs between traits. Moreover, specific reactions were dependent on original as well as target temperature, precipitation and their interactions. It demonstrates the importance of studying these factors simultaneously.

The ecophysiological traits suggest that the populations are able to cope with climate change through plastic response, but it seems that they will probably need genetic adaptation primarily to wetter conditions to maintain their fitness in a long run. We found differences between populations, which could demonstrate a possibility of adaptation to novel conditions, especially in combination with interaction between genotype and phenotype. However, despite a range of significant relationships between ecophysiological and fitness traits, the relationships were quite weak. The strength of the relationships thus need to be very carefully considered in each case to assess biological relevance of the patterns observed.

## Acknowledgements

We would like to thank to H. Skálová for help with setting of physiological measurements, V. Olivová and student helpers for help with data collection and POPEKOL discussion group, T. Herben and P. Mráz for useful comments on the manuscript. This study was supported by project GAČR 19-00522S.

