## Supplementary material for "Ecophysiological traits of a clonal grass in its climate change response"

**Fig. S1.**

The effect of original conditions and target conditions on ecophysiological traits as A) net photosynthetic rate, B) Fv/Fm, C) specific leaf area, D) osmotic potential, E) stomatal density, F) stomatal length. Orig. temp indicates mean summer temperatures at original localities. Orig. moist indicates moisture at the original localities in [mm] of annual precipitation. Columns show mean and error bars +- standard error. Statistical tests are shown in Table 1.

A)

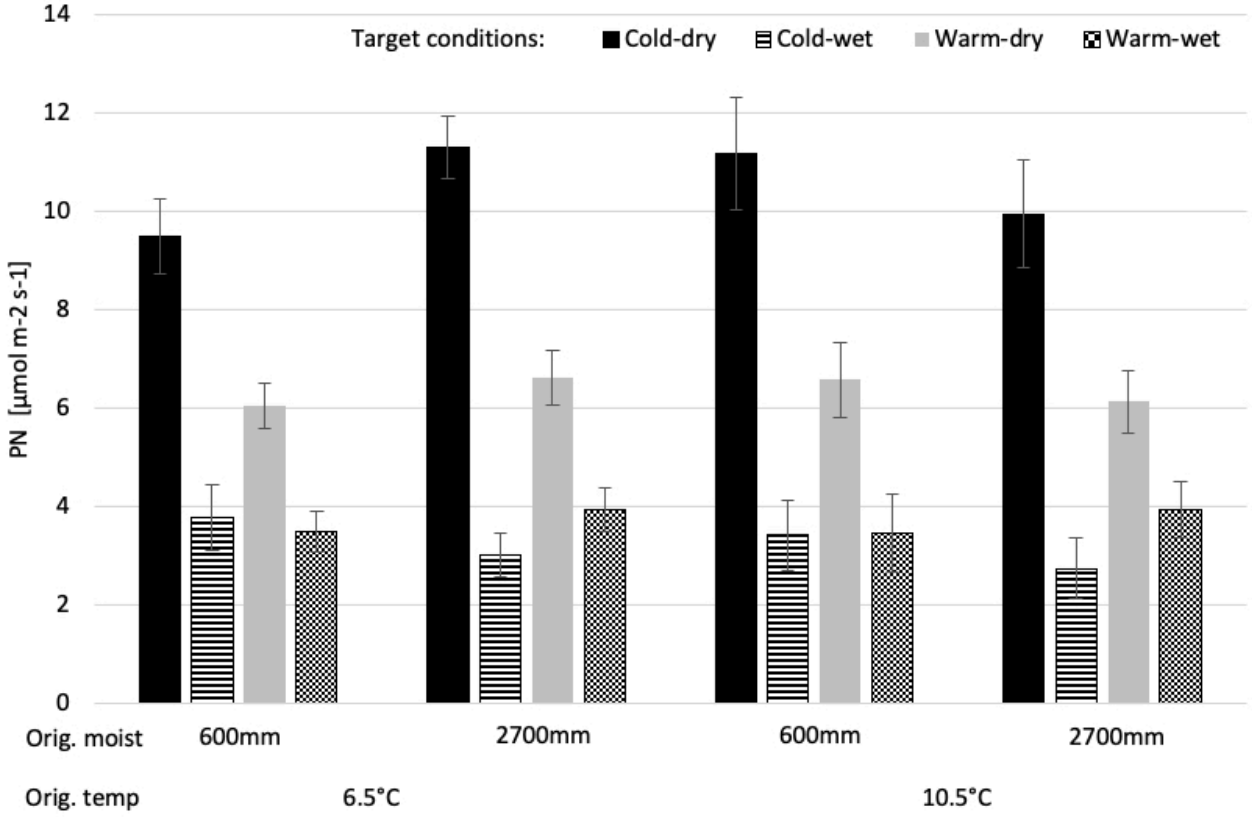

B)

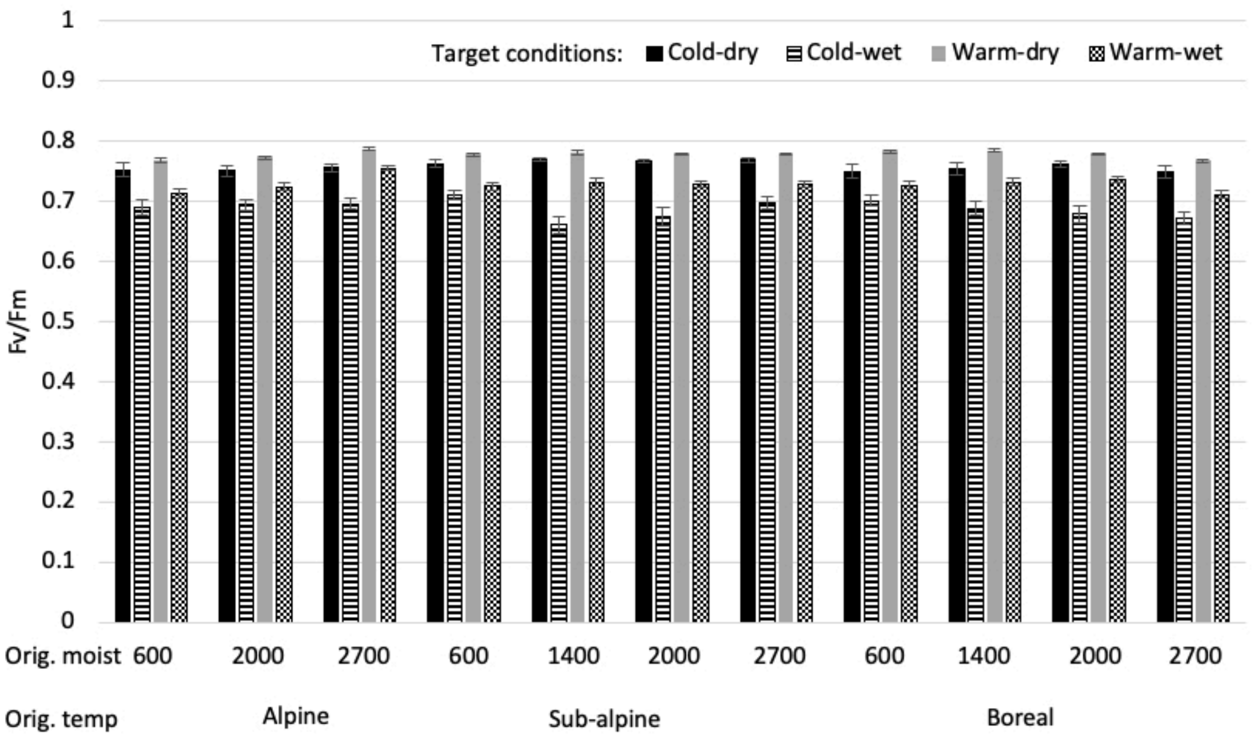
Target conditions: Cold-dry Cold-wet Warm-dry Warm-wet
Fv/Fm
1
0.9
0.8
0.7
0.6
0.5
0.4
0.3
0.2
0.1
0
Orig. moist 600 2000 2700 600 1400 2000 2700 600 1400 2000 2700
Orig. temp Alpine Sub-alpine Boreal

C)

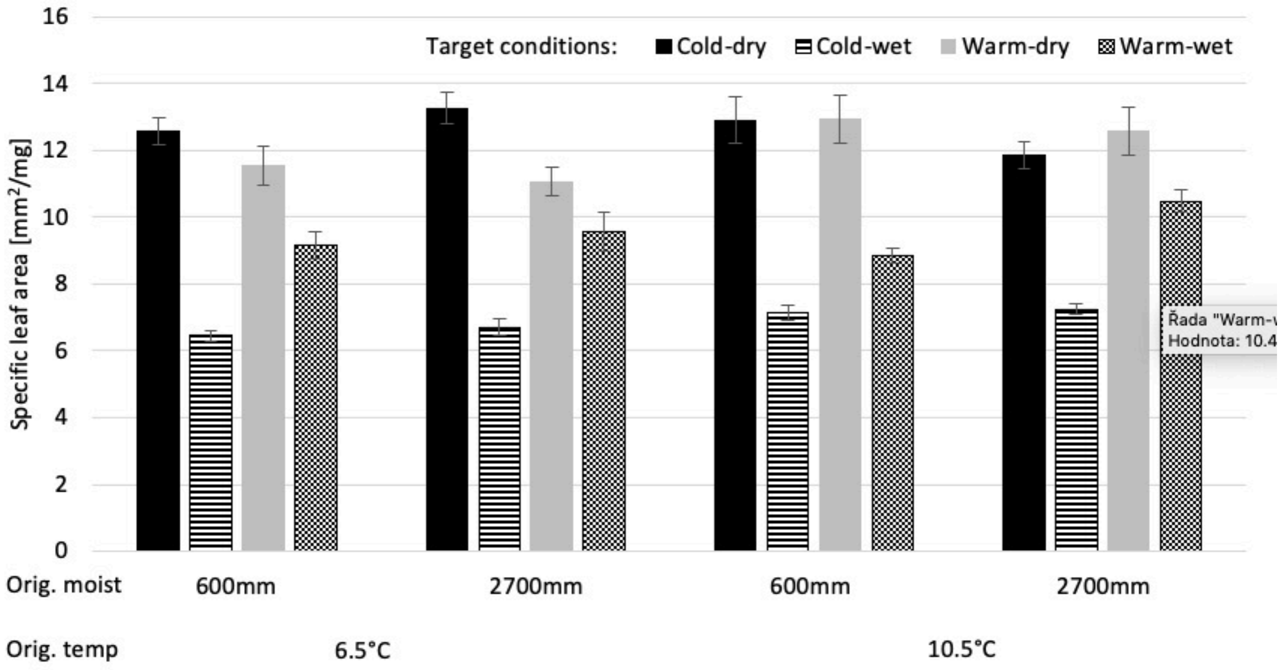
Target conditions:
Cold-dry
Cold-wet
Warm-dry
Warm-wet
Specific leaf area [mm²/mg]
16
14
12
10
8
6
4
2
0
Orig. moist
600mm
2700mm
600mm
2700mm
Orig. temp
6.5°C
10.5°C
Řada "Warm-w
Hodnota: 10.4

D)

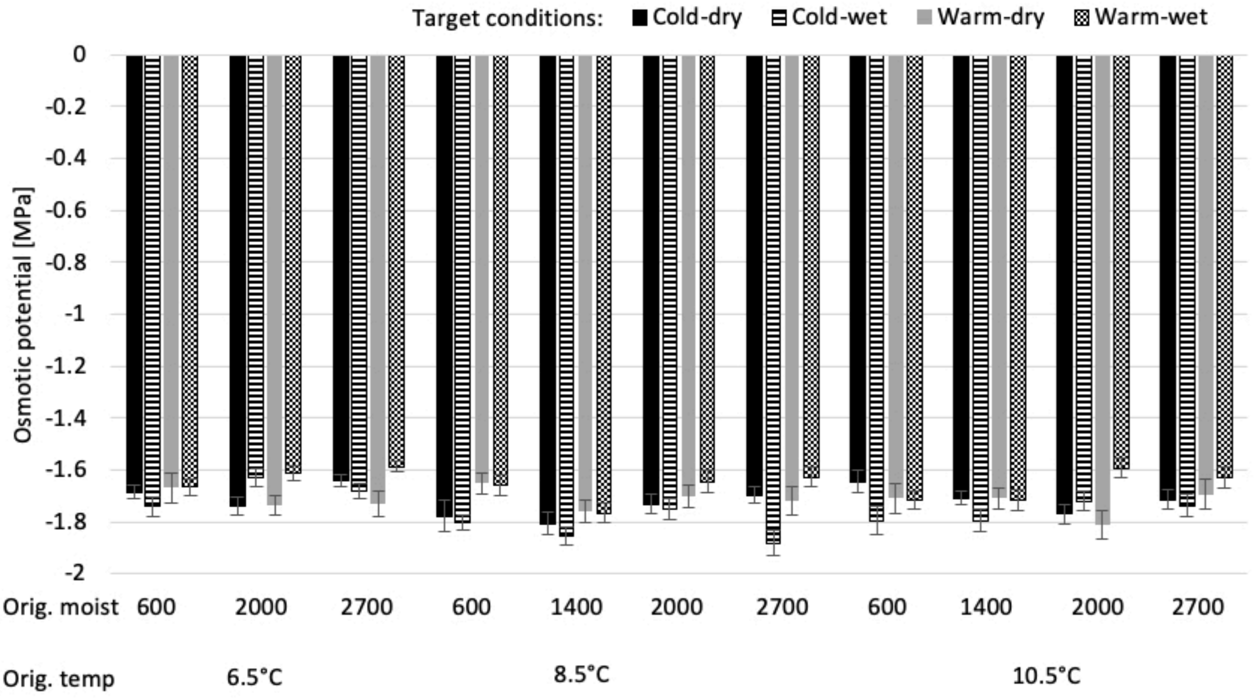
Target conditions:
Cold-dry
Cold-wet
Warm-dry
Warm-wet
Osmotic potential [MPa]
0
-0.2
-0.4
-0.6
-0.8
-1
-1.2
-1.4
-1.6
-1.8
-2
Orig. moist
600
2000
2700
600
1400
2000
2700
600
1400
2000
2700
Orig. temp
6.5°C
8.5°C
10.5°C

E)

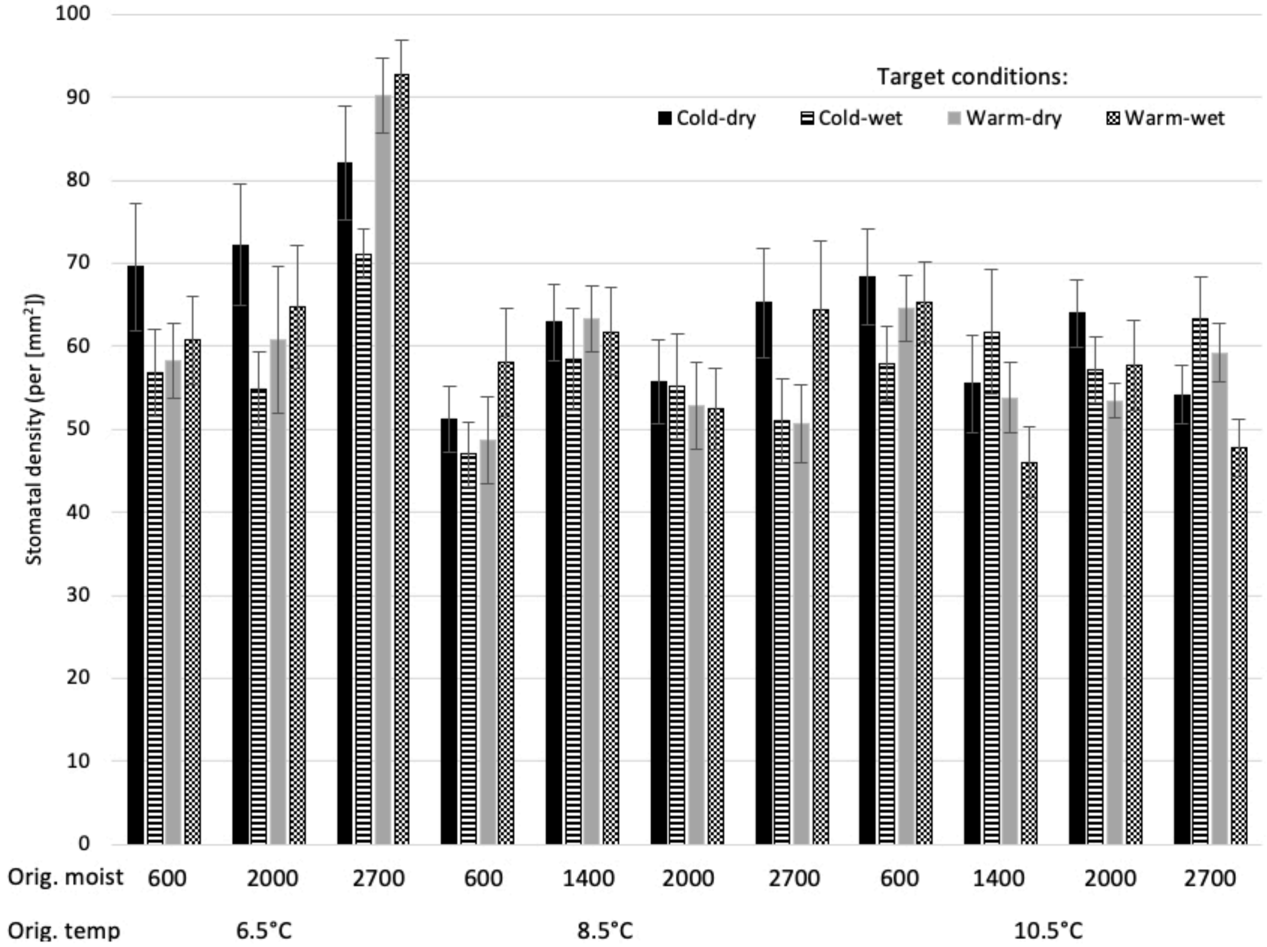
Target conditions:
Cold-dry
Cold-wet
Warm-dry
Warm-wet
Stomatal density (per [mm$^2$])
100
90
80
70
60
50
40
30
20
10
0
Orig. moist
600
2000
2700
600
1400
2000
2700
600
1400
2000
2700
Orig. temp
6.5°C
8.5°C
10.5°C

F)

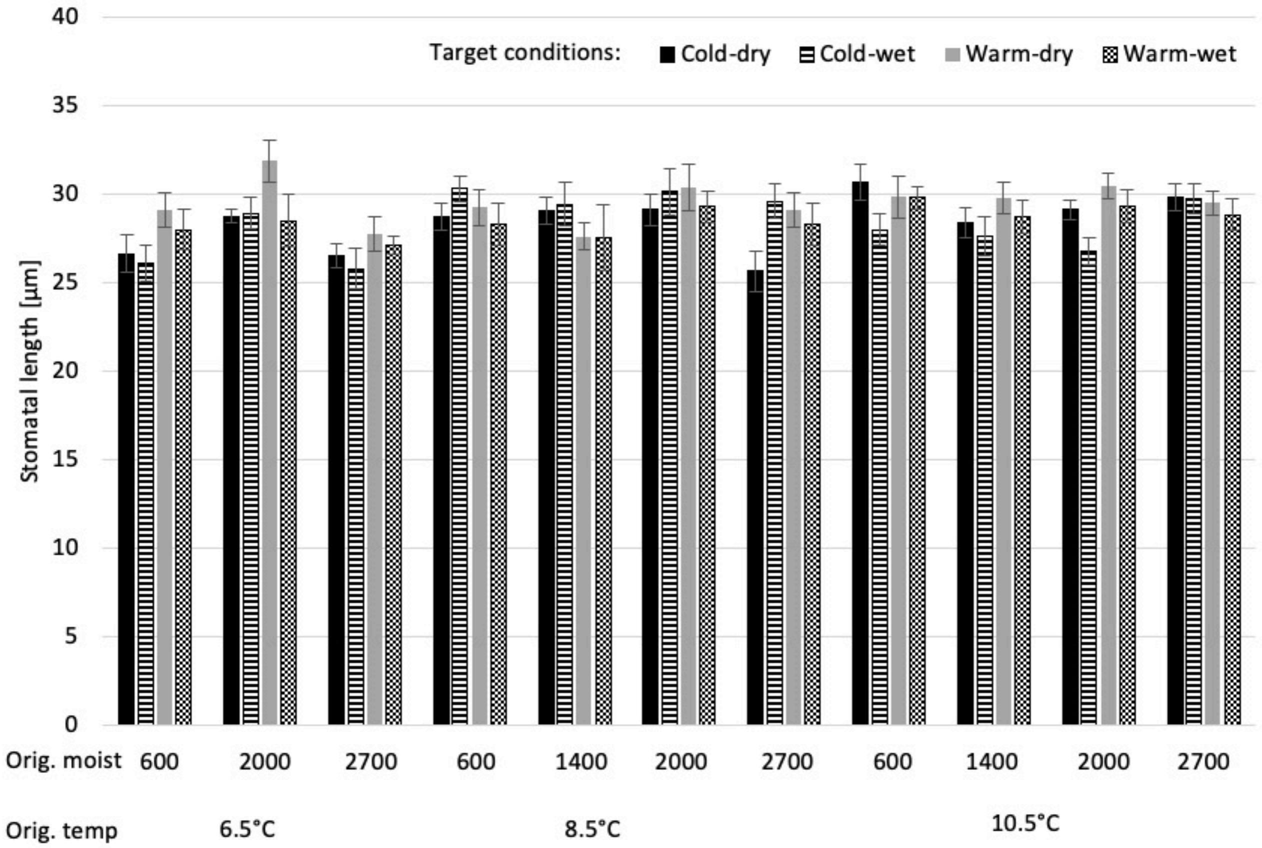
Target conditions:
Cold-dry
Cold-wet
Warm-dry
Warm-wet
Stomatal length [µm]
0
5
10
15
20
25
30
35
40
Orig. moist
600
2000
2700
600
1400
2000
2700
600
1400
2000
2700
Orig. temp
6.5°C
8.5°C
10.5°C

**Fig. S2.**

Effect of difference in temperature and moisture between target and original climate on A) net photosynthetic rate, B) Fv/Fm, C) specific leaf area, D) osmotic potential. The colour scale indicates difference in temperature. Sets of columns indicate differences in moisture. Negative values indicate plants grown in colder and drier conditions, positive values indicate plants grown in warmer and wetter conditions and 0 indicates plants grown in conditions corresponding to the conditions from which they originate. Columns show mean and error bars +- standard error. Statistical tests are shown in Table 2.

A)

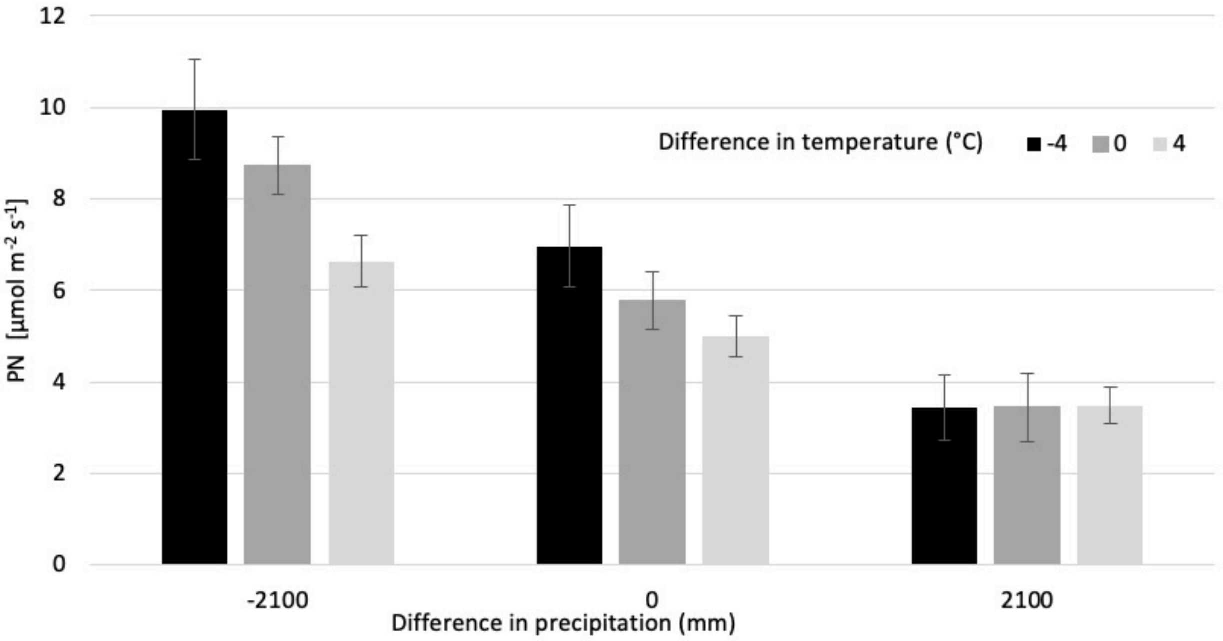

B)

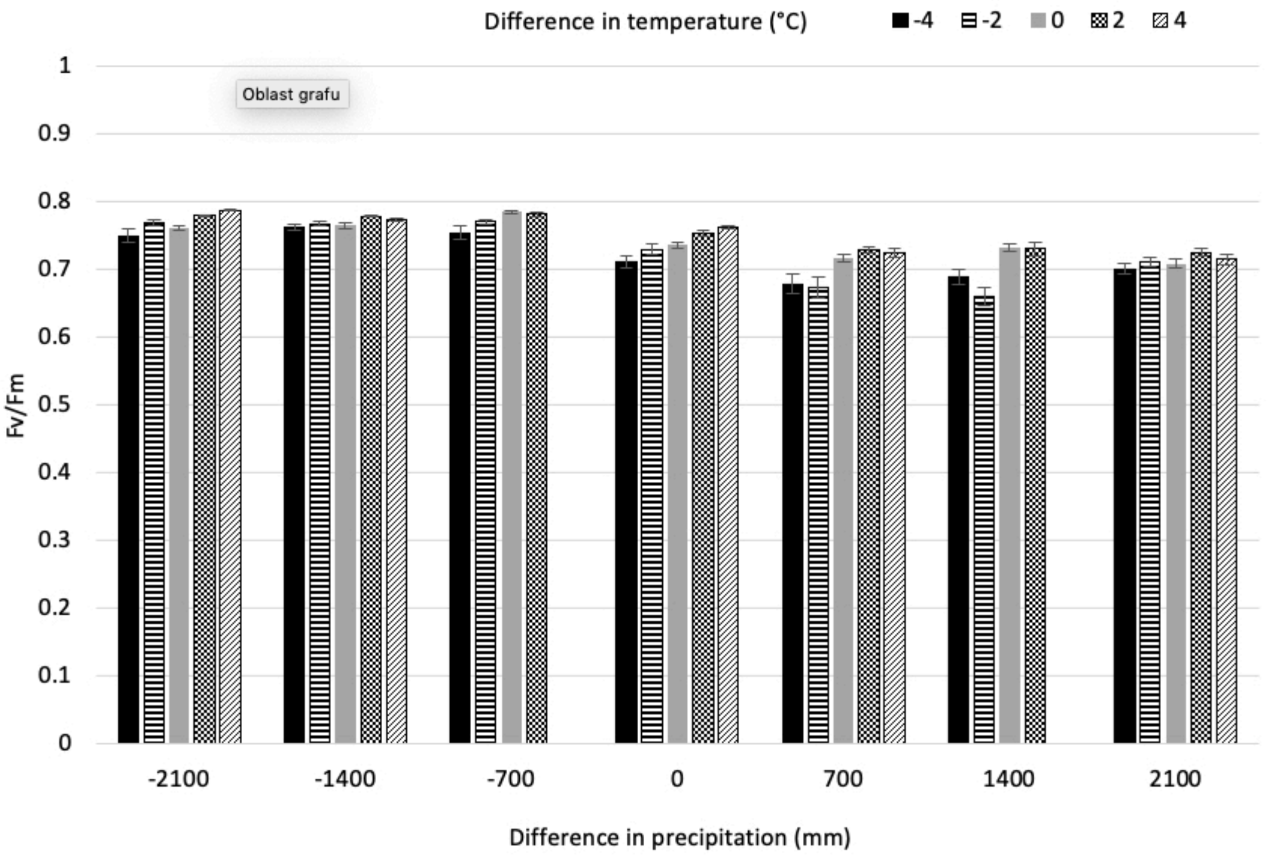

Difference in temperature (°C)
-4
-2
0
2
4
Oblast grafu
1
0.9
0.8
0.7
0.6
0.5
0.4
0.3
0.2
0.1
0
Fv/Fm
-2100
-1400
-700
0
700
1400
2100
Difference in precipitation (mm)

C)

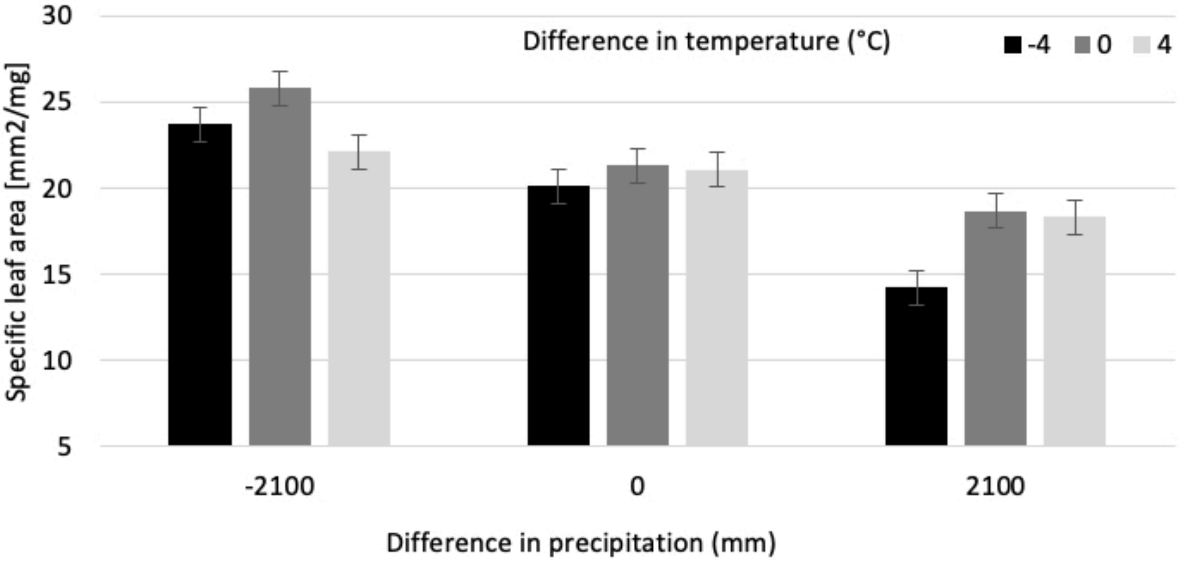

Difference in temperature (°C)
-4
0
4
Specific leaf area [mm2/mg]
30
25
20
15
10
5
-2100
0
2100
Difference in precipitation (mm)

D)

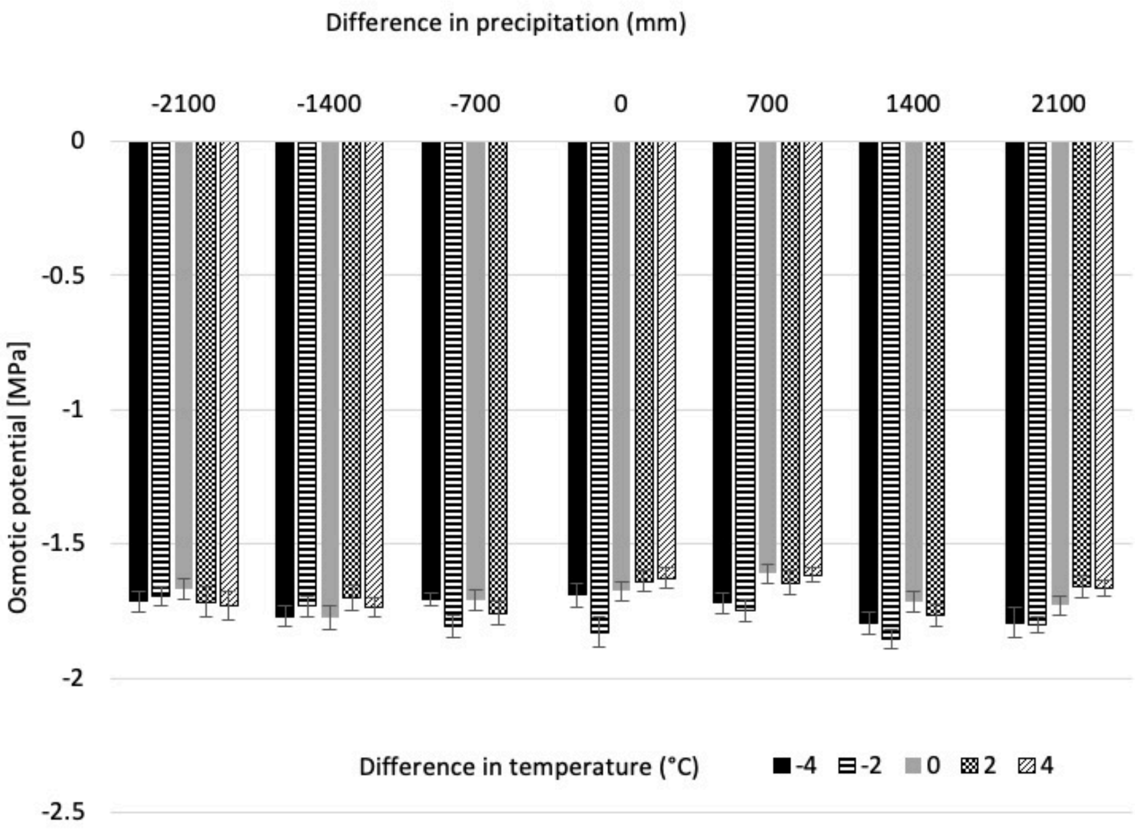

Difference in precipitation (mm)
-2100
-1400
-700
0
700
1400
2100
0
-0.5
-1
-1.5
-2
-2.5
Osmotic potential [MPa]
Difference in temperature (°C)
-4
-2
0
2
4

**Fig. S3.**

A) Effect of local vs. foreign moisture on osmotic potential. B) Effect of interaction between local (L) and foreign (F) temperature (temp) and moisture (mois) on stomatal length. Columns show mean and error bars +- standard error. Statistical tests are shown in Table S3.

A)

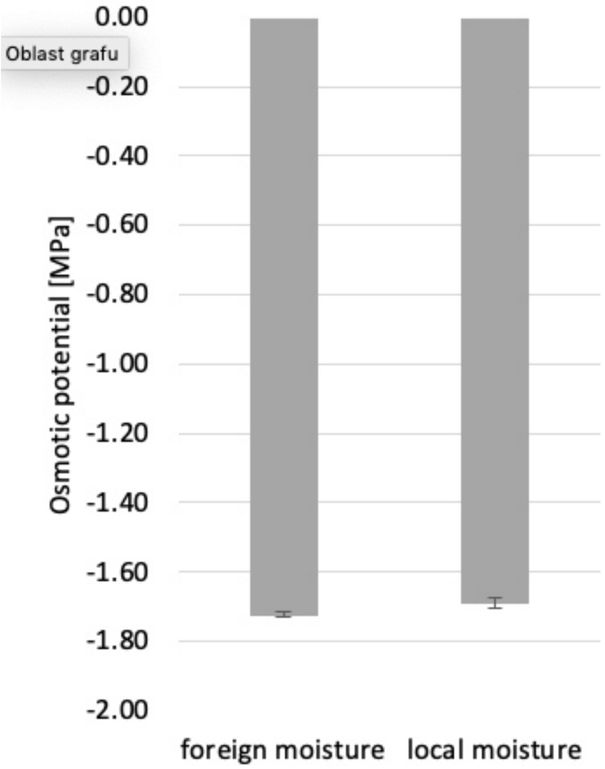

B)

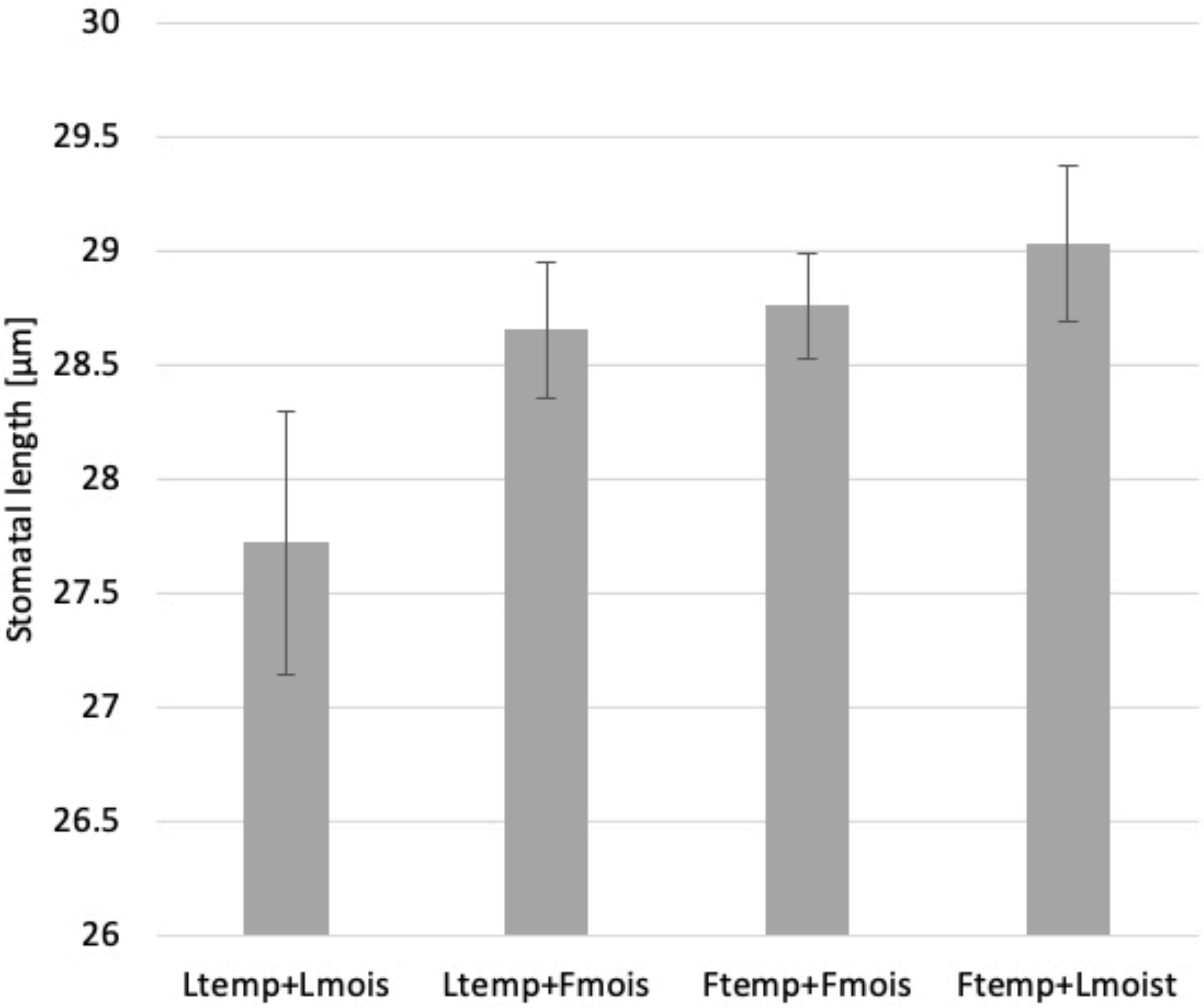
30
29.5
29
28.5
28
27.5
27
26.5
26
Stomatal length [µm]
Ltemp+Lmois
Ltemp+Fmois
Ftemp+Fmois
Ftemp+Lmoist

**Table S1.**
The specific regime settings in the growth chambers providing information on minimum (min), maximum (max) and average (av.) temperature each day.

| Time (day) | Cold regime | | | Warm regime | | |
|---|---|---|---|---|---|---|
| | Min (°C) | Max (°C) | Av (°C) | Min (°C) | Max (°C) | Av (°C) |
| 1-4 | 5 | 15 | 9.8 | 5 | 16 | 10.1 |
| 5-25 | 3 | 12.5 | 7.5 | 3 | 16 | 9.2 |
| 26-46 | 3 | 12.5 | 7.5 | 3 | 18.5 | 10.2 |
| 47-67 | 3 | 12.5 | 7.5 | 3 | 24.3 | 12.5 |
| 68-88 | 3 | 14.5 | 8.4 | 3.4 | 25 | 12.9 |
| 89-176 | 3 | 14.7 | 8.5 | 5 | 23.8 | 14.8 |

**Table S2.**
The effect of original (O) and target (T) temperature (Temp) and moisture (Mois) and their two-way interactions on ecophysiological traits using reduced number of samples (160) for all traits. The tests were done using mixed effect models with genotype as a random factor. Original and target temperature as well as moisture were coded using numbers 1-3 for temperature and 1-4 for moisture, where 1 means the coldest or driest conditions and these numbers were characterized as factor variable. Significant values ($p \leq 0.05$) are in bold. Results marked with * are significant after sequential Bonferoni correction. Dev. indicates deviance explained by the given variable. Net photo. rate ($P_N$) - net photosynthetic rate, Fv/Fm – maximum photosystem II efficiency, *n*– number of observations.

| | Net photo. rate ($P_N$) | | Fv/Fm | | Specific leaf area (SLA) | | Osmotic potential | | Stomatal density | | Stomatal length | |
|---|---|---|---|---|---|---|---|---|---|---|---|---|
| | dev. | p | dev. | p | dev. | p | dev. | p | dev. | p | dev. | p |
| Ttemp | 13.5 | **<0.001*** | 47.74 | **<0.001*** | 7.88 | **0.005*** | 11.26 | **0.001*** | 0.65 | 0.422 | 4.1 | **0.043** |
| Tmois | 118.56 | **<0.001*** | 190.43 | **<0.001*** | 47.32 | **<0.001*** | 21.53 | **<0.001*** | 2.22 | 0.136 | 4 | **0.05** |
| Otemp | 0.27 | 0.81 | 0.43 | 0.514 | 2.97 | 0.085 | 0.44 | 0.506 | 8.5 | **0.004*** | 15.64 | **<0.001*** |
| Omois | 0.01 | 0.722 | 0.45 | 0.501 | 0.32 | 0.573 | 0.07 | 0.786 | 3.48 | 0.062 | 0.17 | 0.677 |
| Otemp:Omois | 0.77 | 0.291 | 7.47 | **0.006*** | 1.08 | 0.3 | 0.86 | 0.353 | 20.75 | **<0.001*** | 0.26 | 0.611 |
| Otemp:Ttemp | 0.01 | 0.824 | 3.75 | 0.053 | 3.3 | 0.07 | 0.01 | 0.96 | 0.08 | 0.16 | 4.78 | **0.029** |
| Otemp:Tmois | 0.29 | 0.851 | 5.85 | **0.016** | 0.82 | 0.364 | 2.55 | 0.11 | 0.08 | 0.776 | 0.04 | 0.842 |
| Omois:Ttemp | 1.18 | 0.453 | 8.78 | **0.003*** | 0.36 | 0.548 | 0.14 | 0.712 | 1.14 | 0.287 | 1.6 | 0.206 |
| Omois:Tmois | 0.1 | 0.995 | 0.19 | 0.005 | 0.01 | 0.96 | 1.23 | 0.268 | 0.28 | 0.6 | 0.47 | 0.493 |
| Ttemp:Tmois | 36.21 | **<0.001*** | 7.97 | **0.005*** | 18.55 | **<0.001*** | 0.15 | 0.695 | 0.75 | 0.385 | 0.24 | 0.627 |
| *n* | 160 | | 160 | | 160 | | 160 | | 160 | | 160 | |

**Table S3.**

The effect of local vs. foreign temperature (Temp) and moisture (Mois) and their two-way interactions on ecophysiological traits. The tests were done using mixed effect models with genotype as a random factor. Significant values ($p \leq 0.05$) are in bold. Results marked with * are significant after sequential Bonferroni correction (for this table, no result is significant after correction). Dev. indicates deviance explained by the given variable. Net photo. rate ($P_N$) - net photosynthetic rate, Fv/Fm – maximum photosystem II efficiency, *n*– number of observations.

| | Net photo. rate ($P_N$) | | Fv/Fm | | Specific leaf area (SLA) | | Osmotic potential | | Stomatal density | | Stomatal length | |
|---|---|---|---|---|---|---|---|---|---|---|---|---|
| | dev. | p-value | dev. | p-value | dev. | p-value | dev. | p-value | dev. | p-value | dev. | p-value |
| Temp | 0.01 | 0.903 | 0.14 | 0.711 | 1.26 | 0.261 | 3.16 | 0.075 | 1.89 | 0.169 | 1.63 | 0.201 |
| Mois | 0.06 | 0.806 | 0.44 | 0.509 | 0.77 | 0.381 | 4.64 | **0.031** | 0.44 | 0.505 | 0.61 | 0.436 |
| Temp:Mois | 0.06 | 0.806 | 1.44 | 0.230 | 0.10 | 0.754 | 0.29 | 0.592 | 0.12 | 0.725 | 4.66 | **0.031** |
| *n* | 160 | | 1100 | | 160 | | 1100 | | 440 | | 440 | |

**Table S4.**

Correlation matrix of physiological traits and growth-related traits. Significant correlation coefficients with $p<0.05$ are marked in bold. Explanatory note: PN – net photosynthetic rate, Fv/Fm – maximum photosysthem II efficiency, SLA – specific leaf area, s. density – stomatal density, s. length – stomatal length, no.ramets – number of ramets, below.biomass – belowground biomass, above.biomass – aboveground biomass

| | PN | Fv/Fm | o.potential | SLA | s.density | s.length | no.ramets | below.biomass | above.biomass |
|---|---|---|---|---|---|---|---|---|---|
| PN | 1 | **0.54** | **0.28** | **0.55** | 0.06 | 0.1 | **0.33** | **0.33** | -0.06 |
| Fv/Fm | **0.54** | 1 | **0.38** | **0.62** | 0.12 | 0.06 | **0.33** | **0.36** | 0.15 |
| o.potential | **0.28** | **0.38** | 1 | **0.28** | -0.12 | 0.15 | 0.06 | 0.14 | 0.05 |
| SLA | **0.55** | **0.62** | **0.28** | 1 | 0.16 | 0.07 | **0.52** | **0.37** | **0.21** |
| s.density | 0.06 | 0.12 | -0.12 | 0.16 | 1 | -0.36 | 0.02 | -0.08 | 0 |
| s.length | 0.1 | 0.06 | 0.15 | 0.07 | -0.36 | 1 | 0.12 | 0.31 | 0.1 |
| no.ramets | **0.33** | **0.33** | 0.06 | **0.52** | 0.02 | 0.12 | 1 | **0.4** | **0.25** |
| below.biomass | **0.33** | **0.36** | 0.14 | **0.37** | -0.08 | 0.31 | **0.4** | 1 | 0.15 |
| above.biomass | -0.06 | 0.15 | 0.05 | **0.21** | 0 | 0.1 | **0.25** | 0.15 | 1 |
